# Direct extrusion of multifascicle prevascularized human skeletal muscle for volumetric muscle loss surgery

**DOI:** 10.1101/2023.09.26.559212

**Authors:** Van Thuy Duong, Thao Thi Dang, Van Phu Le, Thi Huong Le, Chanh Trung Nguyen, Huu Lam Phan, Jongmo Seo, Sung Hoon Back, Kyo-in Koo

## Abstract

Volumetric skeletal muscle injuries are prevalent, highlighting the imperative need for scaffolds to facilitate the healing process of such wounds. Human skeletal muscle is composed of multiple fascicles, which are parallel bundles of muscle fibres surrounded by a layer of connective tissue that contains blood vessels and nerves. Replicating these structures presents a considerable challenge. Here, we developed a method to fabricate multifascicle human skeletal muscle scaffolds that mimic the natural structure of human skeletal muscle bundles using a seven-barrel nozzle. To form the core material to generate the fascicle structure, human skeletal myoblasts were encapsulated in Matrigel with calcium chloride. Meanwhile, to create the shell that plays a role as the connective tissue structure, human fibroblasts and human umbilical vein endothelial cells within a mixture of porcine muscle decellularized extracellular matrix and sodium alginate at a 95:5 ratio was used. We assessed four types of extruded scaffolds monolithic-monoculture (Mo-M), monolithic-coculture (Mo-C), multifascicle-monoculture (Mu-M), and multifascicle-coculture (Mu-C) to determine the structural effect of muscle mimicking scaffold. The Mu-C scaffold demonstrated cell proliferation, differentiation, vascularization, mechanical properties, and functionality that were superior to those of the other scaffolds. Furthermore, in an *in vivo* mouse model of volumetric muscle loss, the Mu-C scaffold effectively regenerated the tibialis anterior muscle defect, demonstrating its potential for volumetric muscle transplantation. The multibarrel nozzle device was applied to create functional Mu-C muscle scaffolds that structurally mimicked human skeletal muscle. Our nozzle will be further used to produce other volumetric functional tissues, such as tendons and peripheral nerves.

## 1 INTRODUCTION

Skeletal muscle is responsible for generating movement forces and accounts for 40–45% of total body mass [1]. Some severe traumatic injuries, such as high-energy fractures, blast injuries, and extensive soft tissue damage or even surgical procedures could cause too volumetric muscle loss to be regenerated by intrinsic muscular regrowth, which leads to scar tissue formation [2]. Unfortunately, this great loss of skeletal muscle results in long-term functional disability [3]. Each skeletal muscle is made up of tens, hundreds, or even thousands of muscle fibres bundled together and sheathed in connective tissues [4]. Each skeletal muscle fibre is a multinucleated muscle cell. On the basement membrane and sarcolemma of each muscle fibre, stem cells differentiate into mature muscle fibres known as satellite cells [5]. A muscle is often divided into groups of muscle fibres called fascicles surrounded by a connective tissue layer called the perimysium. This connective tissue covering provides support and protection for delicate muscle cells [6], allows them to withstand the forces of contraction, and provides pathways for the passage of blood vessels, lymphatics, and nerves.

Muscle transplantation is the gold standard treatment for volumetric skeletal muscle injuries. Free functional muscle transfer involves the use of certain muscles, such as the gracilis muscle or serratus anterior muscle, to restore facial animation during facial muscle surgery [7-9] or to replace the biceps [10]. Even though free functional muscle transfer does not have detrimental functional consequences, it can somewhat adversely affect natural movement, such as flexing the knee, adducting the thigh, and medially rotating the tibia on the femur [11]. The autologous tissues used for implantation often differ in size from the injured muscle, which results in inadequate functional regeneration [12]. To solve this limitation of autologous transfers, donated skeletal muscles that correspond to the injured muscle type can replace the source free functional muscle [13, 14]. However, the lack of available donor tissue [15] and donor site morbidity [14] are two main limitations of donated skeletal muscle transplantation. Given the recent advancements and challenges in obtaining donated sources for skeletal muscle surgery, bioengineered skeletal muscle constructs (BSMCs) have been developed to provide models of mouse muscle [16] or human muscle [17, 18] that are equivalent in terms of physiology and disease to aid in the understanding and treatment of various skeletal muscle disorders and injuries. Currently, two approaches are widely used to generate BSMCs: three-dimensional (3D) casting [16-18] and 3D bioprinting [19-23]. In the 3D casting technique, myoblasts are usually encapsulated in natural hydrogels, such as collagen, Matrigel, or fibrin, to form 3D constructs in a mould with anchors. While this method for the formation of monolithic BSMCs is simple and highly efficient, it is almost impossible to perform inside a mould with multiple materials, such as multifascicle structures. Thanks to using extrusion-based high-resolution nozzles (Ø 200 ∼ 500 µm), the 3D bioprinting approach has the potential to produce multifascicle BSMCs in custom sizes [21]. However, this technique is time-consuming as the high-resolution nozzles can starve and stress cells, leading to necrosis during the printing of high-volume structures [24]. In a study by Choi and colleagues, the highest speed printing process they obtained was calculated to be 0.001 mL/s [21]. Therefore, to construct a muscle with a volume of 10 cm^3^, a minimum of 166.7 minutes would be required. However, this duration is considered excessively long to ensure the survival of cells, particularly when employing multiple cell types for printing biomimetic natural skeletal muscles.

In this study, we propose a method using a multibarrel nozzle to quickly and easily extrude multifascicle skeletal human muscle cells that functionally mimic the natural structure of human skeletal muscle bundles for volumetric muscle loss surgery. Human skeletal myoblasts were expanded *in vitro* and then encapsulated in Matrigel supplemented with calcium chloride (CaCl_2_) as the core material (Figure 1). Human fibroblasts (HFs) and human umbilical vein endothelial cells (HUVECs) covered the myoblast core with a mixture of decellularized extracellular matrix (DECM) and sodium alginate at a ratio of 95:5 as the shell material. Sodium alginate acted as a supporting structure until the gelation of DECM and Matrigel was complete. After complete gelation, the alginate was degraded gradually by alginate lyase. The shell of the extruded scaffolds was similar to the connective tissue layer that covered the entire fascicle. We showed that the HFs and HUVECs within the multi-fascicle structures contributed to forming an interconnected vascular network within the muscle and promoted myogenesis, resulting in higher contractile forces under electrical stimuli and better mechanical properties. Finally, the extruded muscle was matured *in vitro* for 10 days and then implanted as a volumetric muscle to recover the tibialis anterior (TA) muscle defect in mice. These findings would pave the way for the development of an efficient 3D bioprinter capable of constructing volumetric human skeletal muscles.

**Figure 1:**
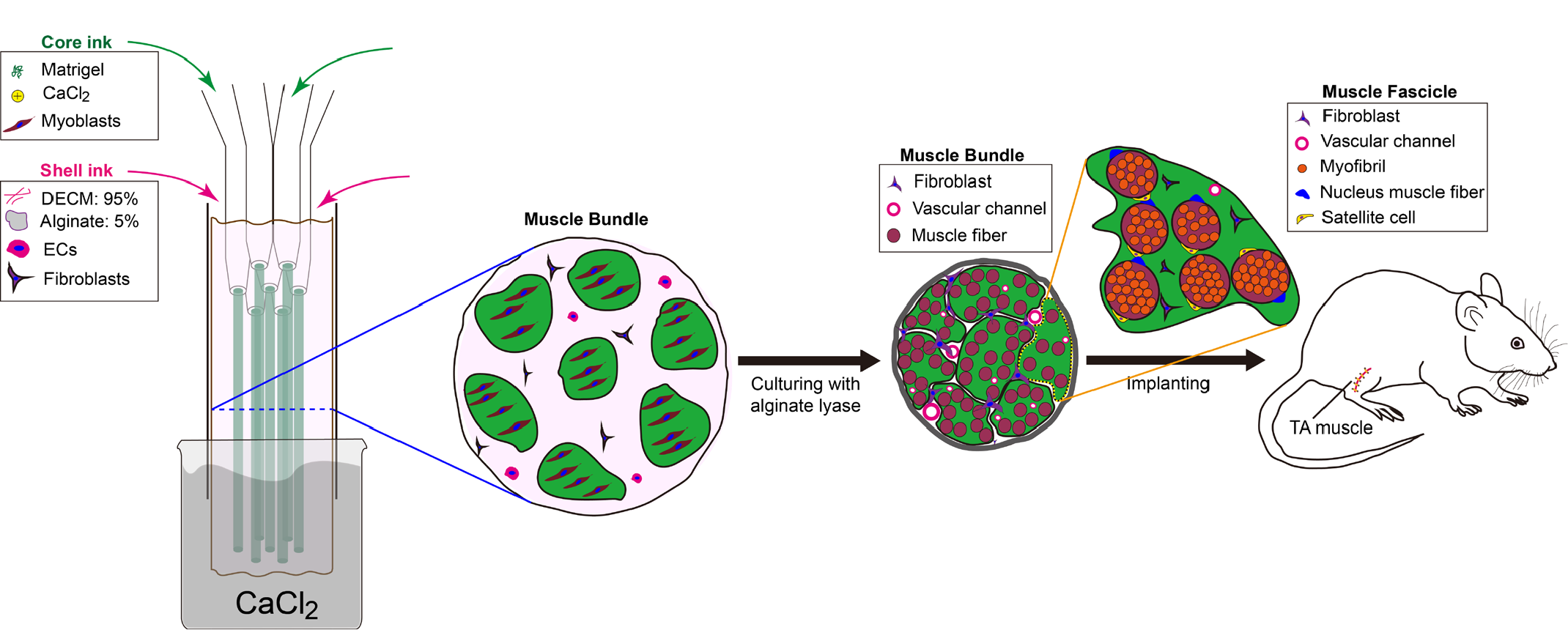
Conceptual representation of multifascicle skeletal muscle extrusion method. The multibarrel nozzle encapsulated human myoblasts within Matrigel as the fascicle material. The larger tube introduced the shell material, which consisted of DECM, sodium alginate, HUVECs, and HFs. The alginate hydrogel was used as a temporary cradle prior to DECM and Matrigel gelation. After gelation, the alginate was dissolved during culture with alginate lyase. HF-HUVEC coculture allowed prevasculature formation and promoted myogenesis during maturation. The extruded muscle was finally implanted into the TA muscle in the right hind limbs of mice.

## 2. RESULTS AND DISCUSSION

### 2.1 ECM-based materials for shell and core

In order to create an optimal microstructure that promotes cell survival, migration, proliferation, and differentiation, we employed ECM-based materials for muscle fabrication. Recently, muscle DECM has been extracted from animals and used as the principal bioink material for tissue fabrication [20, 21, 25, 26]. Muscle DECM contains mainly collagen, laminin, fibronectin, glycosaminoglycans (GAGs), and various growth factors and cytokines, which together create a highly porous structure and promote seeded cells to attach, migrate, proliferate, and differentiate. Figure 2a shows the fibrous structure of our muscle DECM at a 4 wt% concentration; this structure contained fibril matrices with submicrometer diameters. These fibrous structures play a vital role in supporting cell function, adhesion, migration, and signalling, contributing to tissue development, maintenance, and repair [27]. After gelation, the muscle DECM hydrogels were very soft and difficult to manipulate as they broke during handling (Figure S1a). Indeed, the storage modulus (G’) of the muscle DECM hydrogels created by Hernandez *et al*. was measured to be approximately 5 Pa at 4 wt%, 10 Pa at 6 wt%, and 17 Pa at 8 wt% [28]. The gelation time for muscle DECM under physiological conditions (37 °C and pH 7) has been demonstrated to range from 10 to 30 min [20, 21], which is unsuitable for an extrusion-based bioprinter. Therefore, to increase the durability and reduce the gelation time of extruded hydrogels, we mixed muscle DECM and sodium alginate at a ratio of 95 to 5 as the shell material. This hydrogel mixture still contained microfibrils and porous structures (Figure 2b). Compared with pure DECM, the mixed DECM exhibited better durability during handling in experiments (Figure S1). Moreover, the bioink mixture stiffened and retained its shape after being extruded from the outlet and reacting with CaCl_2_ in a harvesting container (Video S1). Hence, the combination of 95% DECM and 5% alginate proved to be an appropriate formulation for the scaffold’s shell.

**Figure 2:**
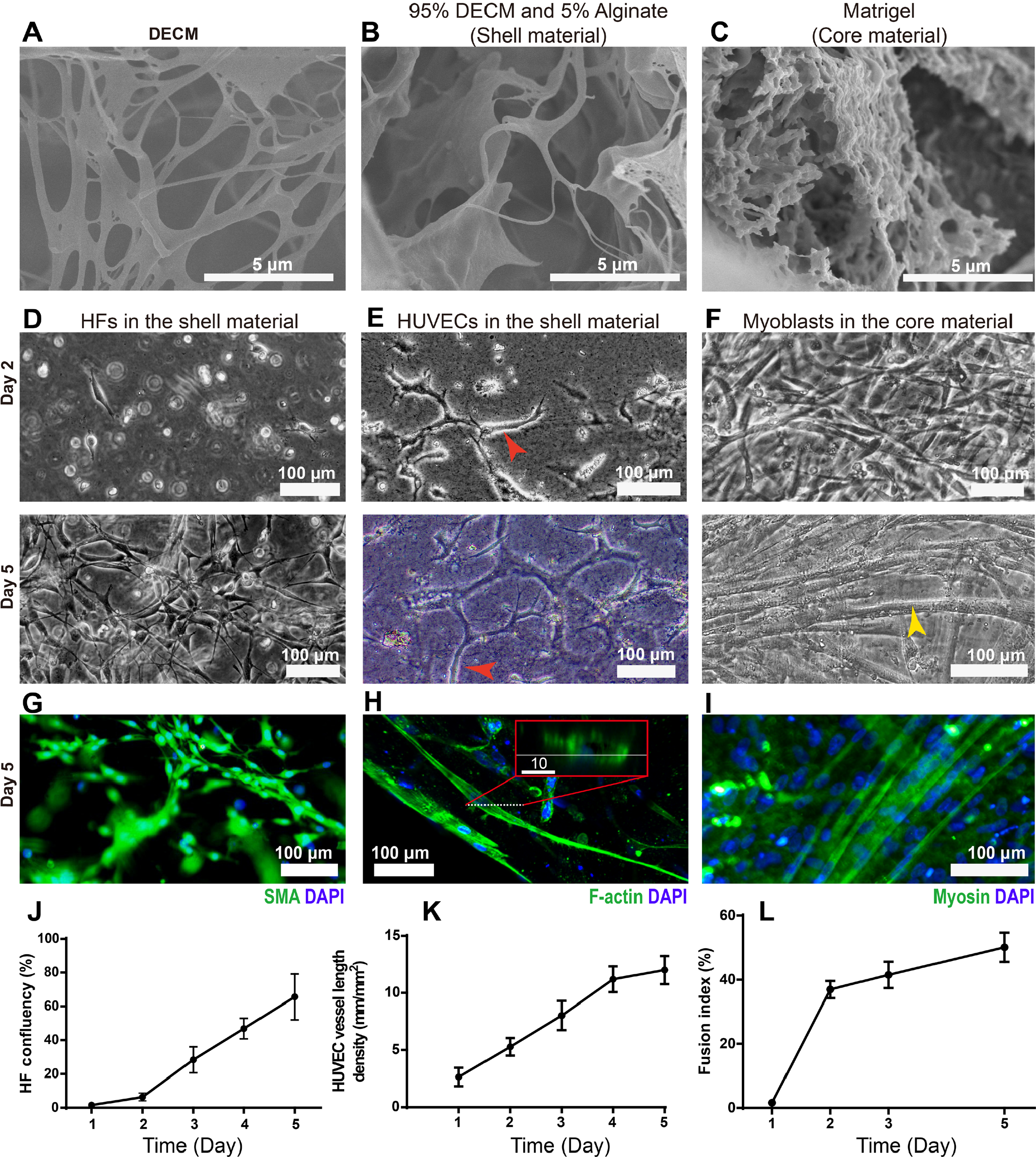
The microstructures of. (a) DECM, (b) 95% DECM −5% sodium alginate (shell material), and (c) Matrigel (core material). (d) HFs and (e) HUVECs were cultured in the shell hydrogel. The red arrows indicate microvascular vessels. (f) Human myoblasts were cultured in the core hydrogel. The yellow arrow indicates a typical muscle fibre differentiated from myoblasts. Immunofluorescence staining of (g) HFs and (h) HUVECs in the shell material. (i) Immunofluorescence staining of human myoblasts in the core material. (j) HF confluency in the shell hydrogel (n = 3). (k) Quantification of the HUVEC vessel length density (n = 4). (l) Quantification of the fusion index of human myoblasts (n = 4).

Of note, the calcium in the core material is also involved in crosslinking the shell material prior to extrusion from the outlet [29]. The characteristics of our materials demonstrated their printability. Figure 2c shows the microstructure of Matrigel, which contained very small cavities approximately 0.5 to 3 µm in size. Due to this micro structure and many other molecular cues contained in Matrigel, this material offers various advantages, such as cytocompatibility, biomimicry to natural ECM, and the capacity to promote cell attachment, migration, and differentiation across many cell types [30]. Therefore, in this investigation, Matrigel was used as the core material for human skeletal muscle derived cells (hSkMDCs) to form muscle fascicles.

HFs, HUVECs, and hSkMDCs were cultured in bulk gels made of shell and core materials to examine their confluence, morphology, and viability for 5 days. Figure 2d-f and Figure S2 show bright field images of HFs, HUVECs, and hSkMDCs in the shell and core materials. HF confluence was approximately 8-fold higher on day 5 than day 2 (Figure 2d, g, j). The HUVECs exhibited network-like morphology on day 2 (Figure 2e, h). A lumen structure was observed in the network-like morphology on day 5 (Figure 2h, Figure S3). Particularly, the total length of the HUVECs in one square millimetre on day 5 was approximately two times longer than that on day 2 (Figure 2k). This tube formation by HUVECs is a fundamental biological process that plays a crucial role in the development of blood vessels within engineered skeletal muscles [31]. The viability of the HFs and HUVECs ranged from 92% to 95% throughout the whole culture period (Figure S4). These results indicate that our shell material provides a favourable environment for HFs and HUVECs to proliferate, migrate, and exert their biological functions. hSkMDCs proliferated quickly in 2 days (Figure 2f). Once differentiated, they migrated and merged to form multinucleated cells within the core hydrogel (Figure 2f, i, Figure S5) and exhibited robust and spontaneous contractions by day 5 (Video S2). This transformation is vital for muscle function, development, repair, and growth, enabling efficient contractile activity and adaptation to various physiological demands. Specifically, by utilizing the 3D casting technique, Juhas *et al*. demonstrated that the multinucleated cell transformation of human myoblasts formed structurally and functionally biomimetic skeletal muscle [16].

To evaluate the ratio of differentiated hSkMDCs within the core material, *Choi*’s fusion index was utilized [20]. The fusion index in the core material revealed that approximately 55% of the hSkMDCs had differentiated in 5 days (Figure 2l). This index was equivalent to the result reported by Choi et al. for human myoblast differentiation within 3D bioprinted constructs [20].

### 2.2 Multi-fascicle muscle structure

In order to create multi-fascicle scaffold, we developed a method to fabricate multifascicle human skeletal muscle scaffolds that mimic the natural structure of human skeletal muscle bundles using a seven-barrel nozzle and evaluated the method by comparing the properties of four extruded models and their performance in vivo. Figure 3a-c shows the fabricated nozzle and extrusion setup and process. A cell-free 7 fascicle scaffold was smoothly extruded into a beaker containing CaCl_2_ (Figure 3c, Video S1). Food dyes were used to distinguish the fascicles from the shell. We also demonstrated the printability of our material by extruding a 7-fascicle scaffold on a glass slide (Video S1). When extruded on the glass slide, a moderate concentration of calcium cations (25 mM) in the core material stimulated crosslinking with the sodium alginate in the shell material, resulting in gradual gelation at the outlet. This gelation maintained the three-dimensional shape of the extruded scaffold on the slide during printing.

**Figure 3:**
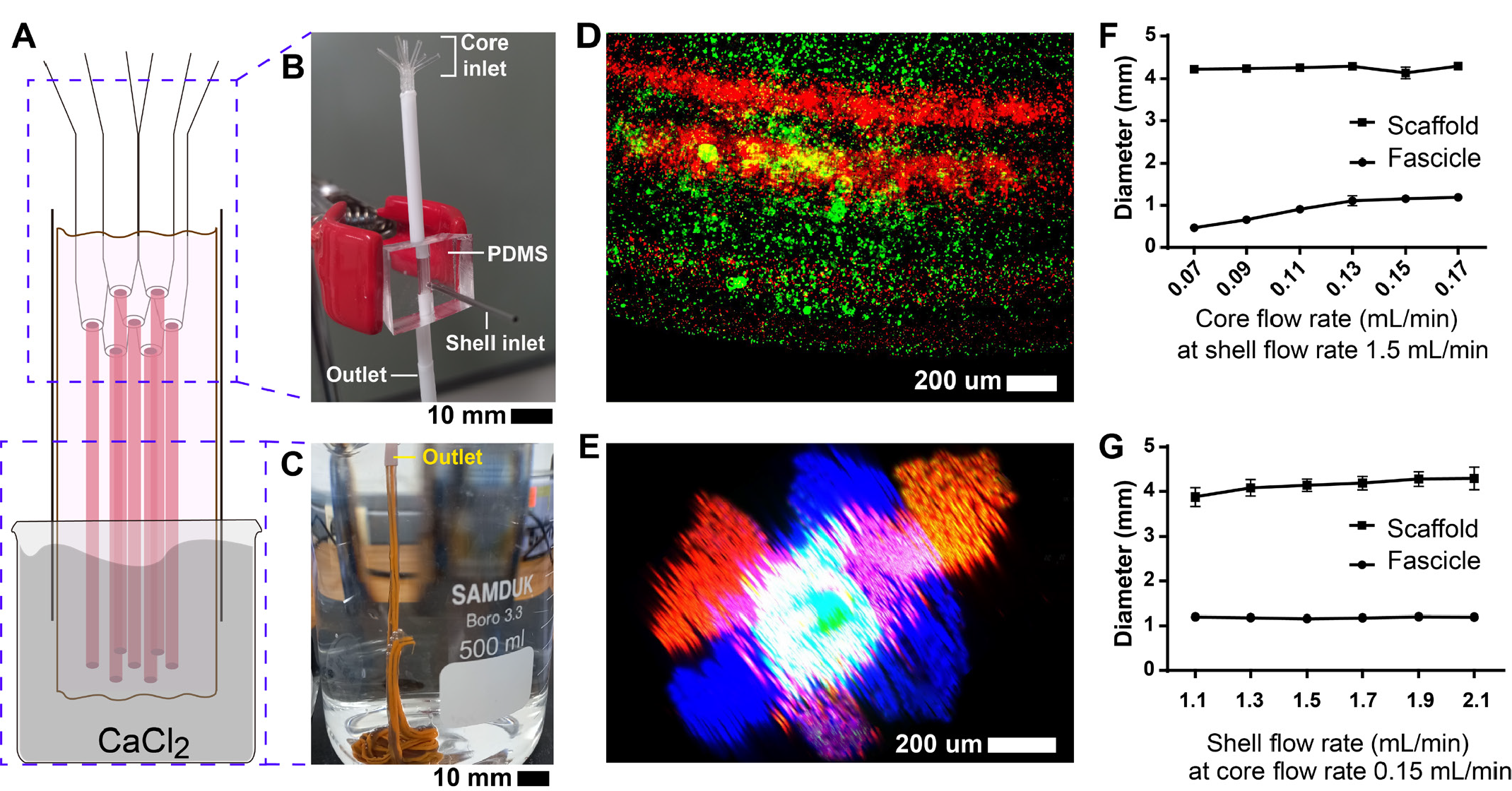
Cell-free scaffold extrusion with the proposed multibarrel nozzle device. (a) Schematic of the multifascicle extruding device. (b) The fabricated extruding device. (c) Cell-free multifascicle scaffold extrusion into a bath of CaCl_2_. (d) The multifascicle scaffold observed under a fluorescence microscope. (e) Cross-section of the 7-fascicle scaffold. Red and blue were used to distinguish fascicles. (f) The change in scaffold diameter as a function of shell flow rate (n = 5). (g) The change in scaffold diameter as a function of core flow rate (n = 5).

To examine the feasibility of bioprinting with the multifascicle device, fluorescent microparticles (PS-FluoGreen-Fi199, S-FluoRed-Fi198, and S-FluoBlue-Fi165; microParticles GmbH, Germany) were mixed with the shell and core materials and extruded. Subsequently, fluorescence images were acquired that displayed the multifascicle structure of the extruded scaffold (Figure 3d, e). The circular cross-section of the scaffold showed an asterisk structure with seven interspersed fascicles. The fascicle and scaffold diameters could be controlled by changing the input flow rate. Increasing the core flow rate from 0.07 to 0.17 mL/min at a fixed shell flow rate of 1.5 mL/min increased the fascicle diameter from 0.76 to 1.2 mm, while the outer diameter of the scaffold remained approximately 4.2 mm (Figure 3f). On the other hand, the scaffold outer diameter increased from 3.8 to 4.3 mm, and the fascicle diameter remained approximately 1.2 mm when the shell flow rate increased from 1.1 to 2.1 mL/min at a constant core flow rate of 0.015 mL/min (Figure 3g). Thus, fascicle diameter was more sensitive to variation in the core flow rate than in the shell flow rate. The fascicle diameter of our scaffold, approximately 1 mm, is similar to the fascicle diameter of mammalian muscle [32]. Therefore, our proposed device was able to create multi-fascicle scaffolds structurally mimicking natural skeletal muscle.

Once we succeeded with cell-free scaffold extruding, we proceeded the extrusion with cell encapsulation. Figure 4 shows the extruded multifascicle scaffolds cells e on days 0, 2, and 10. The scaffold was relatively fragile immediately after extrusion but became more durable after 2 days and could be easily handled after 10 days. Fascicles within the scaffold could be observed from the outside as white stripes along the scaffolds which were visible to the naked eye 2 days after differentiation.

**Figure 4:**
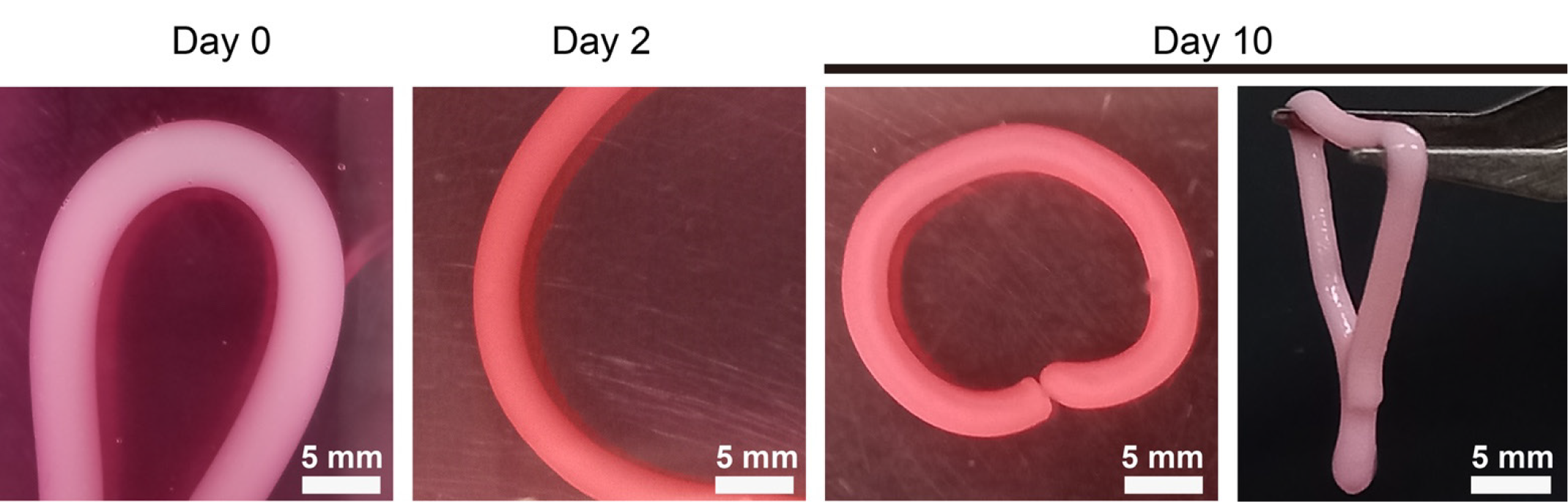
The generated, cultured cell-laden scaffold. The multifascicle cell-laden scaffold was relatively fragile immediately after extrusion but became increasingly durable starting on the second day and durable enough to handle starting on the tenth day.

Under confocal microscopy, these stripes exhibited signals indicating immunostained Myosin (green in Video S3). This result indicated that the encapsulated hSkMDCs differentiated and generated contractile units known as primary motor protein responsible for converting chemical energy stored in adenosine triphosphate (ATP) into mechanical work. This process enables muscles to contract and generate force [16].

To evaluate the multifascicle structure with multiple cells, four types of muscle scaffolds were formulated: monolithic–monoculture (Mo-M), monolithic–coculture (Mo-C), multifascicle–monoculture (Mu-M), and multifascicle–coculture (Mu-C) (Figure 5a). The two multifascicle scaffolds (Mu-M and Mu-C) were generated by the device proposed in this investigation (Figure 3), and the two monolithic scaffolds (Mo-M and Mo-C) were formulated by our previously developed device for two-layered blood vessels [29]. Cross-sections of the extruded muscle scaffolds on day 10 exhibited unique structural features (Figure 5b). Using F-actin staining, it was easy to distinguish the multifascicle scaffolds with cross-sectional asterisk shapes. All muscle scaffolds shrank approximately 30% in size 2 days after extrusion, and their diameters were maintained in the range of 2.1 mm to 2.4 mm (Figure 5c). The Mu-C scaffold displayed the biggest reduction in size without significance. This shrinkage caused the scaffold compact and denser with cells (Figure 4).

**Figure 5:**
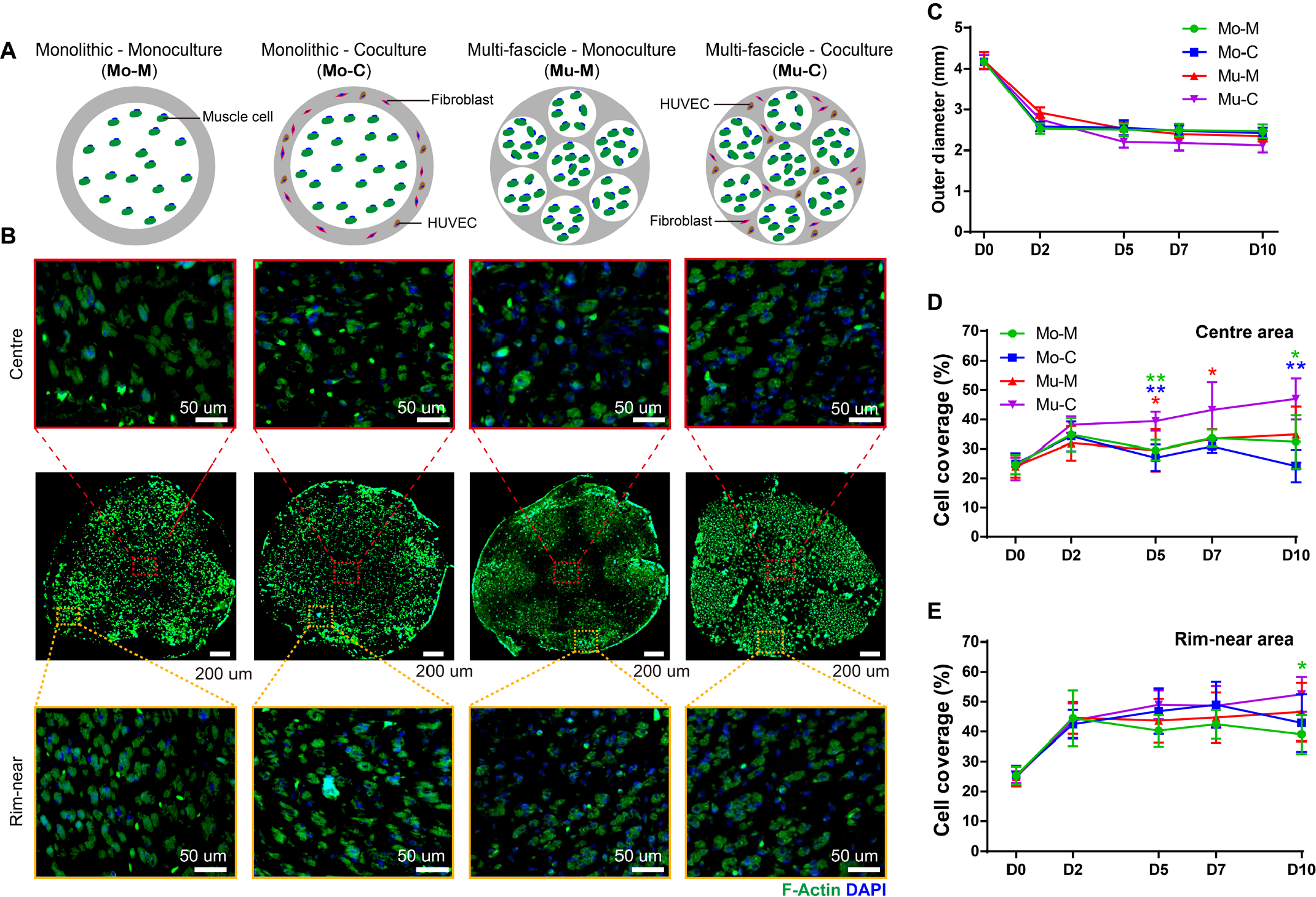
Four types of cell-laden scaffolds. (a) Conceptual schematics of differences in the cell-laden scaffolds. (b) Cross-sections of the cell-laden scaffolds. The sections were immunostained with F-actin and DAPI. The first and third rows are higher magnifications of the centre row. (c) The change in outer diameter of the cell-laden scaffolds according to culture time (n = 6). (d) The cell coverage ratio in the centre area of the cell-laden scaffolds according to culture time (* 0.01 < p < 0.05, ** 0.001 < p < 0.01 comparing the Mu-C group to the other groups; n = 4). (e) The cell coverage ratio at the peripheral area of the cell-laden scaffolds according to culture time (* 0.01 < p < 0.05, comparing the Mu-C group to the other groups; n = 4).

Cell coverage on the cross-section of the cultured muscle scaffolds was quantified for 10 days (Figure 5d and e). In the first two days, the encapsulated cells quickly increased their coverage at both the centre and peripheral areas in all the muscle scaffolds. The cells in the centre area in the Mu-C scaffold displayed significantly greater coverage than those of the other scaffolds beginning on day 2. The multifascicle structure and coculture were expected to facilitate the development of a microvascular network around the muscle fascicles. This enhancement would facilitate the diffusion of nutrients, thereby promoting the viability and proliferation of the enclosed cells [33]. Compared with Mu-M, fibroblasts and HUVECs appeared to enhance the proliferation of centre muscle cells. Overall, the cell coverage in the peripheral area was approximately 10% higher than that in the centre area. It was therefore suggested that the cells in the peripheral area could more feasibly diffuse in cell culture media than those in the centre area.

### 2.3 Myogenesis

Myogenin (MyoG) is a muscle-specific basic helix-loop-helix transcription factor involved in the coordination of skeletal muscle development or myogenesis and regeneration [16, 34]. To show the differentiation of myoblasts into mature muscle fibers, regulating various stages of myogenesis within the muscle structures, we immunostained horizontally cross-sectional slices with myogenin and DAPI. At the peripheral area, hSkMDCs differentiated quickly, as 40%, 62%, and 80% of the cells expressed MyoG on days 5, 7, and 10, respectively (Figure 6a, b). At the early stage of muscle formation (day 2), approximately 10% and 6% of cells were positive for MyoG at the peripheral and centre areas of all muscle scaffolds, respectively (Figure 6b). There was no large difference in MyoG expression at the peripheral areas among the 4 types of muscle scaffolds over 10 days. However, in the centre area of the Mu-C muscle scaffold, there were many more MyoG-positive hSkMDCs on days 5, 7, and 10 than on the other muscle scaffolds (Figure 6c, d). This result demonstrated that the multifascicle structure and coculture provided favourable conditions for hSkMDC differentiation. Similarly, by culture myoblasts for day 14 within Matrigel scaffolds, Mark Juhas *et al*. also demonstrated that about 80% of the cells differentiated into myofibers and expressed MyoG [16].

**Figure 6:**
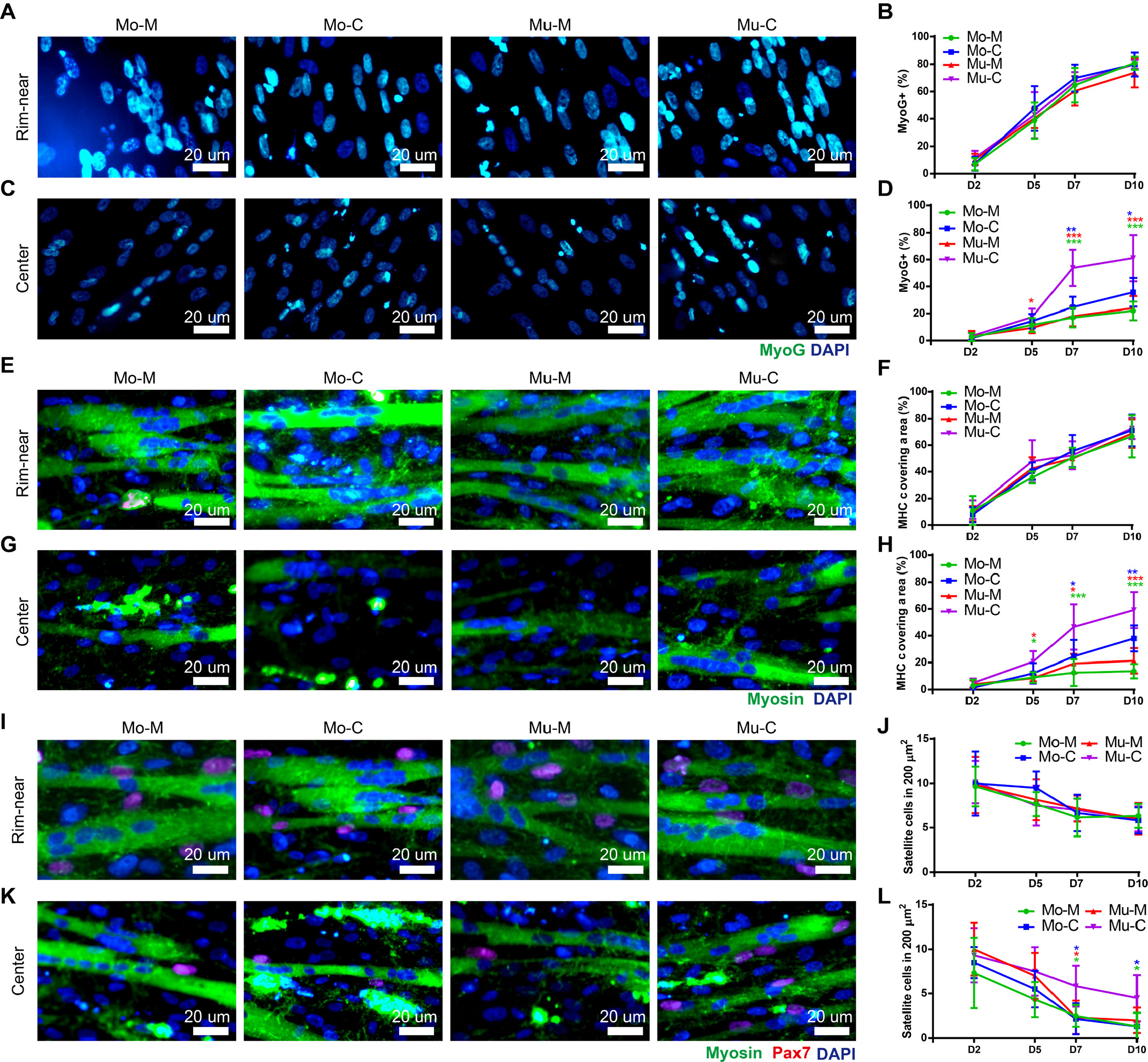
Myogenesis of myoblasts within fascicles of the scaffolds. (a) Immunofluorescence detected Myogenin and DAPI on day 10 in the peripheral areas. (b) Percentage of Myogenin-positive nuclei at the peripheral areas (n = 6) over time. (c) Immunofluorescence of Myogenin on day 10 in the centre areas. (d) Percentage of Myogenin-positive nuclei in the centre areas as a function of culture time (* 0.01 < p < 0.05, ** 0.001 < p < 0.01, ***p < 0.001 comparing the Mu-C group to the other groups; n = 6). (e) Immunofluorescence of Myosin on day 10 in the peripheral areas. (f) Percentage of Myosin coverage in the peripheral areas as a function of culture time (n = 6). (g) Immunofluorescence of Myosin at day 10 in the centre areas. (h) Percentage of Myosin coverage in the centre areas as a function of culture time (* 0.01 < p < 0.05, ** 0.001 < p < 0.01, ***p < 0.001 comparing the Mu-C group to the other groups; n = 6). (i) Images of muscle fibres coimmunostained with myosin, Pax7, and DAPI on day 10 in the peripheral areas. (j) Number of satellite cells in the peripheral areas as a function of culture time (n ≥ 6). (k) Images of muscle fibres coimmunostained with myosin, Pax7, and DAPI on day 10 in the centre areas. (l) Number of satellite cells in the centre areas as a function of culture time (* 0.01 < p < 0.05, n ≥ 6).

Next, we immunostained the muscle scaffolds with Myosin antibodies to show multinucleated muscle fibres as myosins are a superfamily of motor proteins known for their roles in muscle contraction [35]. The coverage of Myosin-positive cells illustrated cell fusion to form muscle fibres [16]. On day 10, the Mu-C muscle scaffold had significantly higher Myosin coverage (approximately 60%) than the other scaffolds. However, the Myosin coverage in the peripheral area increased without any significant difference among all the scaffolds, even on day 10 (Figure 6e, f). Especially on day 10, the Mu-C muscle scaffold at the peripheral area exhibited approximately 65% Myosin coverage, which was not significantly different from that at the centre area. At the centre area, the Myosin+ hSkMDC coverage of all the scaffolds showed an increase with increasing culture time (Figure 6 g, h). These results indicated that hSkMDCs within the Mu-C muscle differentiated more quickly and evenly than those within the other muscles.

Pax7 is considered a marker protein of satellite cells, and when it is highly expressed, it usually means that satellite cells are activated and can proliferate to form new muscle fibres [36]. Therefore, to determine the density of satellite cells, Pax7 and Myosin were coimmunostained. The average densities of the satellite cells in both areas were 8.79 cells in 200 µm^2^, without any significant difference during the early stages of muscle formation (day 2) (Figure 6i-l). The density decreased significantly on days 5, 7, and 10. The densities of the satellite cells in the peripheral areas were significantly higher than those in the centre areas of Mo-M, Mo-C, and Mu-M muscles beginning on day 2. However, there was no significant difference between the peripheral and centre areas of the Mu-C muscle. This result suggested that the multifascicle structure and coculture provided suitable conditions to promote progenitor development for muscle regeneration.

Of note, the muscle fibres at the centre area of the Mu-C muscle were significantly larger than those of the other muscle scaffolds on days 7 and 10 (Figure 7a). The diameters of the muscle fibres in the Myosin immunostained images increased from approximately 10 µm on day 2 to approximately 20∼25 µm on day 10 in the peripheral area of all the muscle scaffolds (Figure 7b). These diameters were equivalent to those of human native muscle fibres (> 18 µm) [37]. The muscle fibres in the centre areas were thinner than those in the peripheral areas in all the muscle scaffolds.

**Figure 7:**
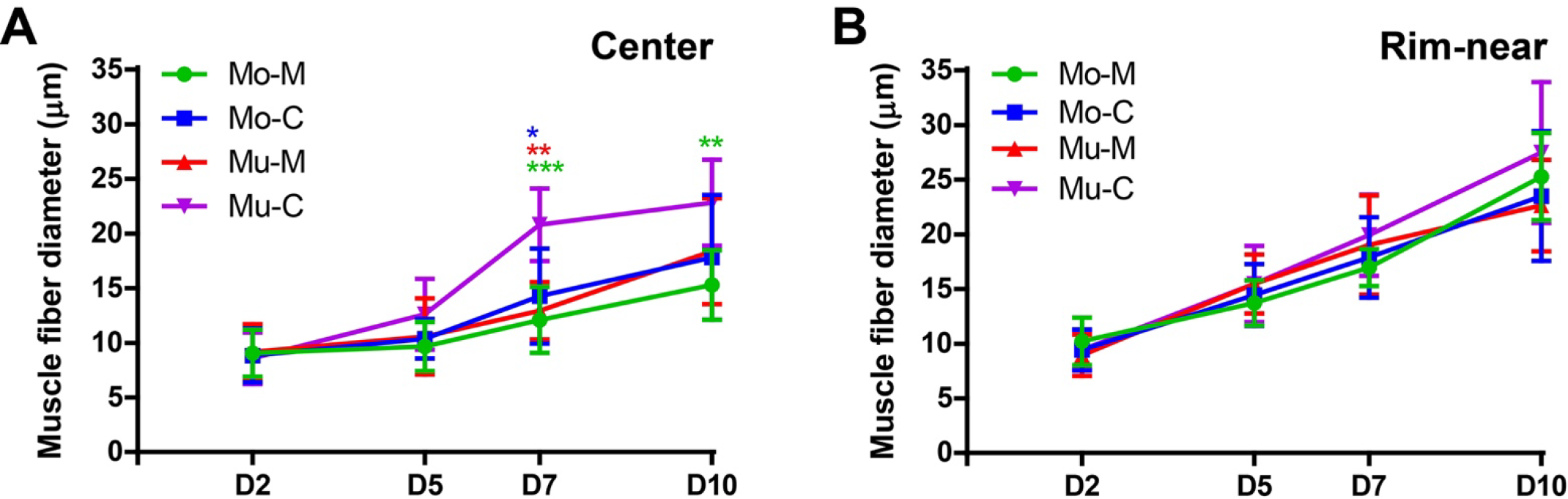
Muscle fibre diameter. (a) The change in average myofiber diameter in the centre areas as a function of culture time (* 0.01 < p < 0.05, ** 0.001 < p < 0.01, ***p < 0.001 comparing the Mu-C group to the other groups; n ≥ 6). (b) The change in average myofiber diameter at the peripheral areas as a function of culture time (n ≥ 6).

### 2.7 Contractile force after electrical stimulation

We measured the contractile force when the fibres were stimulated with electrical pulses from a function generator to assess the muscle functionality of our multifascicle scaffolds. The Mu-C scaffold exhibited a significantly stronger force than all the other scaffolds under all stimulation conditions (Figure 8). All the muscle scaffolds contracted more intensively at a stimulation frequency of 50 Hz stimuli than at 10 Hz (Figure 8a, c). In the comparison of the contractile force at an electrical stimulation frequency of 50 Hz on day 5, all muscles contracted more intensely at 25 volts/cm stimulation than at 15 volts/cm, and the Mu-C scaffold exhibited significantly higher contractile forces than the other muscle scaffolds (Figure 8b). All the muscle scaffolds (Mo-M, Mo-C, Mu-M, and Mu-C) showed stronger contraction on day 10 when stimulated with 25 volts/cm (Figure 8c) and 50 volts/cm (Figure 8d, Video S4, Video S5). The highest contractile force (20 to 25 mN) was generated by the Mu-C scaffold at 25 volts/cm and 50 Hz of stimulation on day 10. These results indicated that the higher cell proliferation and myogenesis of hSkMDCs within the Mu-C scaffold increased its contractile force. This functional muscle scaffold could be applied as an implantable muscle *in vivo* or as a model of muscle physiology and disease to aid in the treatment of various muscle disorders.

**Figure 8:**
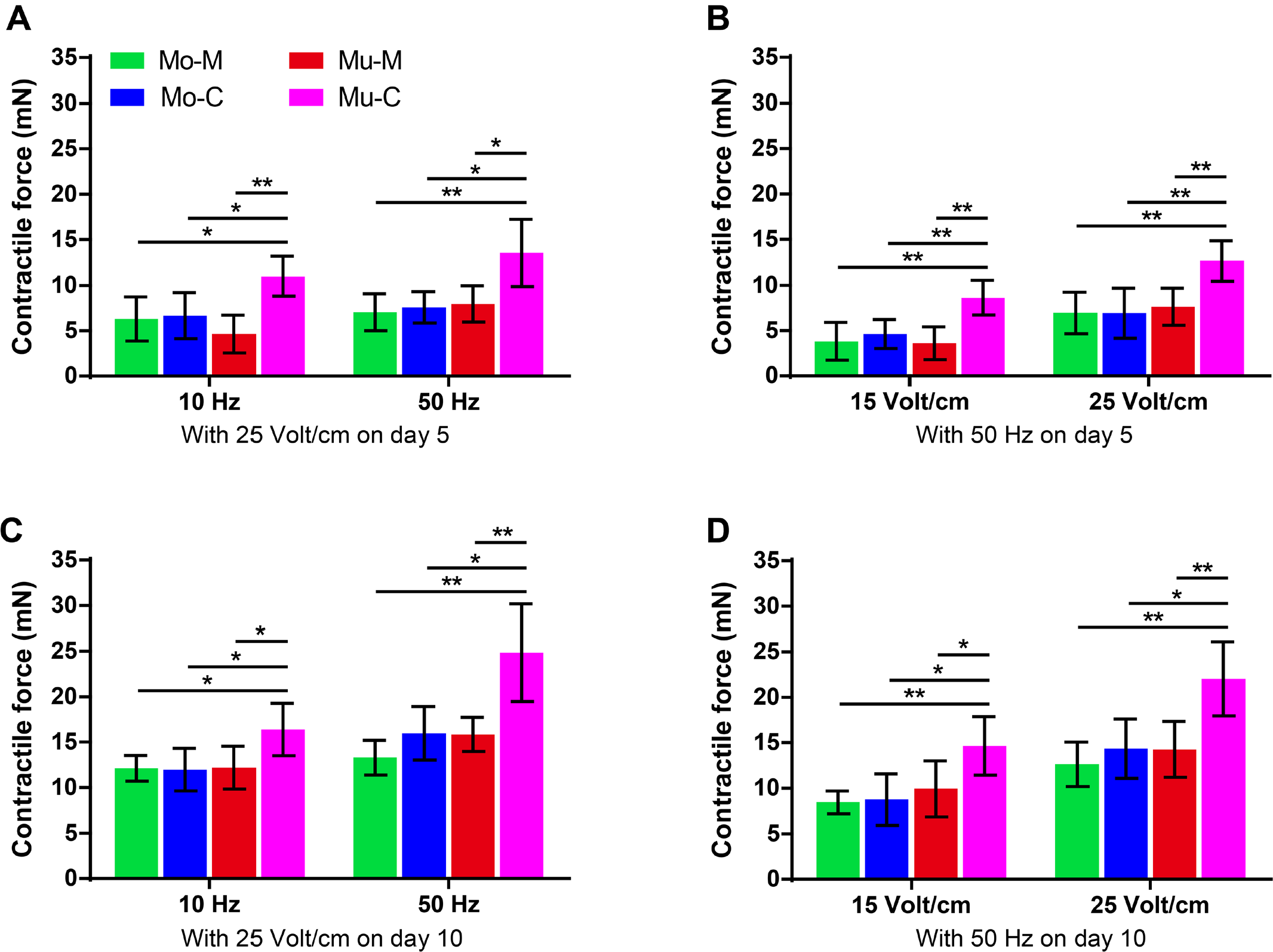
Contraction under electrical stimulation. (a) Contractile forces on day 5 at two different stimulus frequencies (* 0.01 < p < 0.05, ** 0.001 < p < 0.01 comparing the Mu-C group to the other groups; n = 5). (b) Contractile forces on day 5 at two different stimulus voltages (** 0.001 < p < 0.01 comparing the Mu-C group to the other groups; n = 5). (c) Contractile forces on day 10 at two different stimulus frequencies (* 0.01 < p < 0.05, ** 0.001 < p < 0.01 comparing the Mu-C group to the other groups; n = 5). (d) Contractile forces on day 10 at two different stimulus voltages (* 0.01 < p < 0.05, ** 0.001 < p < 0.01 comparing the Mu-C group to the other groups; n = 5).

### 2.5 Vascular network formation

Samples were immunostained with CD31 to show the presence of endothelial cells within the muscle scaffolds. Figure 9a, b, c shows the histological immunofluorescence of CD31 within the coculture muscle scaffolds (Mo-C and Mu-C) on day 5 at the peripheral, centre, and shell areas (the grey area in Figure 5a). There was no significant difference between the peripheral and shell areas with respect to HUVEC coverage on day 5 in both types of muscle scaffolds (Figure 9a, b, f). However, HUVEC coverage at the centre area of the Mu-C scaffold was significantly higher than that of the Mo-C scaffold (Figure 9c, f). This result indicated a denser microvascular network within the Mu-C muscle than in the other scaffolds.

**Figure 9:**
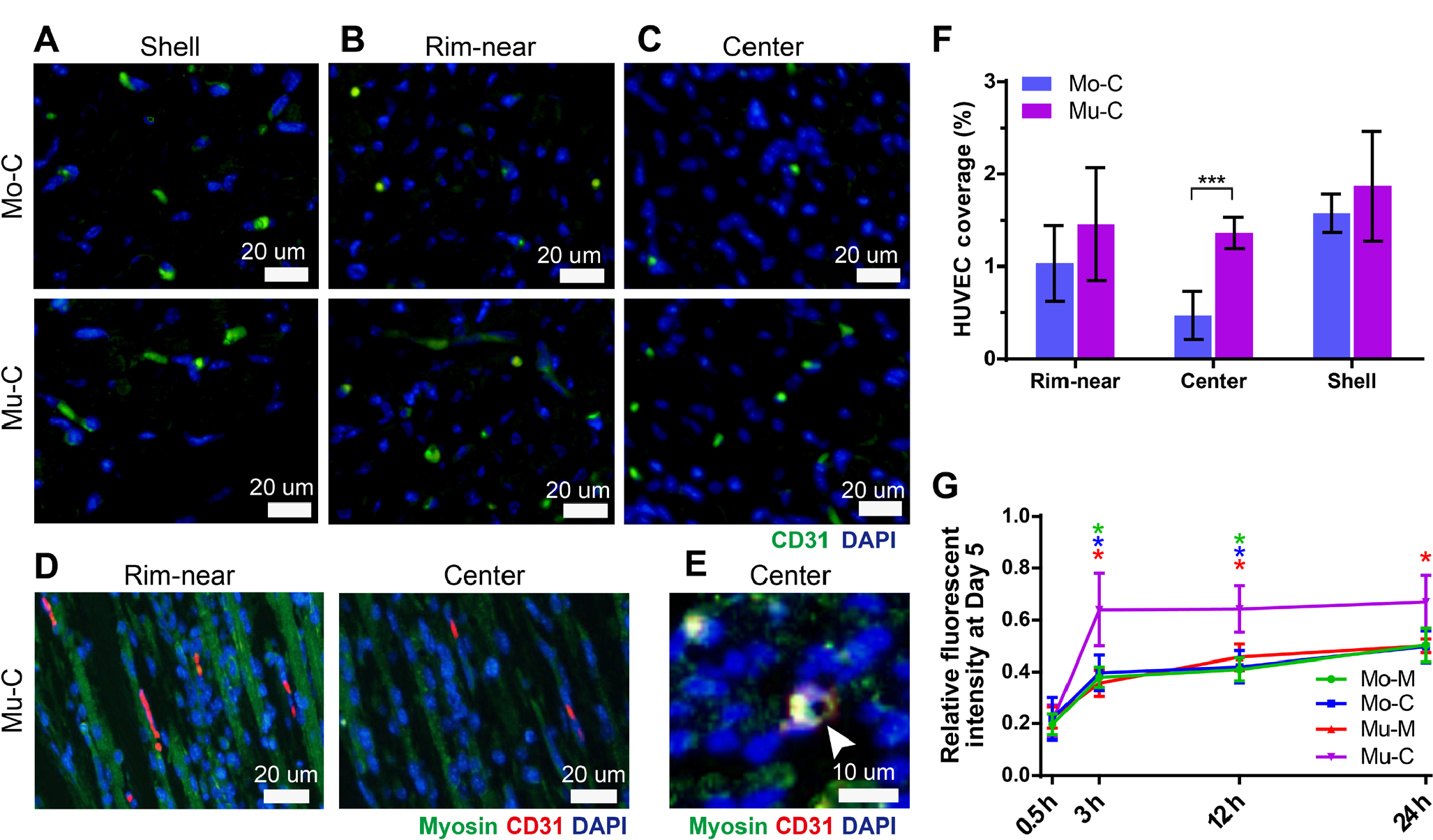
Vascularization within muscle scaffolds. CD31 immunofluorescence images of the cocultured scaffolds on day 5 in the (a) shell areas, (b) peripheral areas, and (c) centre areas. (d) Immunofluorescence images of the Mu-C scaffolds labelled with Myosin, CD31, and DAPI. (e) The lumen structure (white arrow) in the immunofluorescence image of the Mu-C scaffold labelled with Myosin, CD31, and DAPI. (f) Percentage of HUVEC coverage according to position (***p < 0.001, n ≥ 6). (g) Relative fluorescence intensity as a function of time after the scaffold was immersed in 70 kDa FITC-dextran to demonstrate the diffusability of scaffolds (* 0.01 < p < 0.05, comparing the Mu-C group to the other groups; n = 3).

In the longitudinal section images, the interconnecting vascular channels look to be intertwined with the muscle fibres (Figure 9d). The lumen structure of a microvascular channel at the centre area of the Mu-C scaffold was observed, the diameter of the channel was approximately 4 µm, and it was surrounded by CD31 signals (Figure 9e).

On day 5, identical amounts of the 4 muscle scaffolds were immersed in fluorescent 70 kDa dextran, and the relative fluorescence intensity was measured in digested slurry solution (Figure 9g) to find out the difference in dextran diffusability between the scaffolds. The Mu-C scaffold showed significantly higher diffusion of dextran than the other scaffolds after 3 hours. The interconnecting microvascular network was expected to increase dextran diffusion, resulting in a higher relative fluorescence intensity. This result implied that the denser microvascular network interconnected the shell, peripheral, and centre areas to enhance nutrient diffusion from the surrounding milieu outside the Mu-C scaffold into deeper layers. This superior diffusion seemed to promote the hSkMDCs within the Mu-C scaffold to better proliferate and differentiate into functional muscle fibres (Figure 5 and Figure 6).

This assumption was also partially demonstrated by Choi et al. through using a 3D cell printed muscle construct containing microvascular channel [21]. Our vascularizing strategy in this study might be an efficient approach to enhance vascular density and homogeneity leading to an increment in the diffusion distance of nutrients and oxygen to deeper layers of volumetric muscle constructs [38].

### 2.6 Durability

To assess the durability of muscle scaffolds, a mechanical measurement system was established and utilized for 20-mm muscle samples, as illustrated in Figure 10a. In the stress−strain curves of the 4 types of muscle scaffolds on day 10, at the same level of strain, the Mu-C scaffold required higher stress than the others (Figure 10b). Figure 10c shows the change in ultimate stress of all the scaffolds over 10 days. All the muscles were very soft and fragile on day 2, but their durability increased approximately 15-fold in three days. The Mu-C scaffold showed significantly higher ultimate stresses on days 5, 7, and 10 compared with the other scaffolds (VIdeo S6).

**Figure 10:**
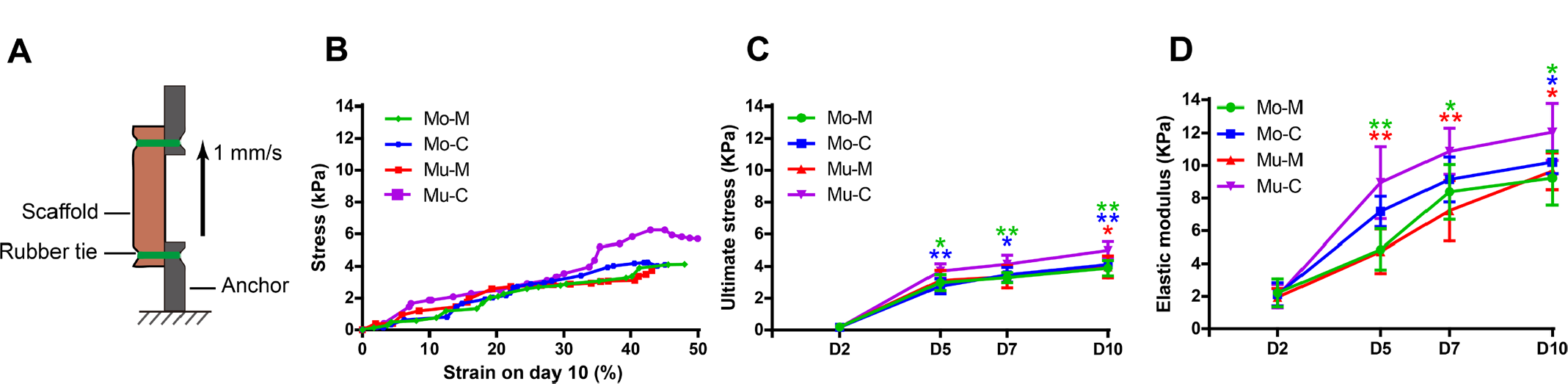
Mechanical properties of the extruded cell-laden scaffolds. (a) Conceptual schematic of the tensile force measuring system. (b) The stress−strain curves of the extruded cell-laden scaffolds on day 10. (c) The ultimate stress as a function of culture time (* 0.01 < p < 0.05, ** 0.001 < p < 0.01 comparing the Mu-C group to the other groups; n ≥ 5). (d) The elastic modulus as a function of culture time (* 0.01 < p < 0.05, ** 0.001 < p < 0.01 comparing the Mu-C group to the other groups; n ≥ 5).

Based on the ultimate strain and stress, the elastic moduli were calculated and plotted in Figure 10d. There were no significant differences in elastic moduli among the groups on day 2. However, after day 5, the elastic modulus of the Mu-C scaffold was significantly higher than that of the other scaffolds. On day 10, the Mu-C scaffold exhibited an elastic modulus of approximately 11.85 kPa, which was close to that of native muscle tissue (10∼12 kPa) [39, 40]. These results demonstrated that the multifascicle and coculture conditions provided the hSkMDCs with a much more suitable structure so that they could exhibit native muscle tissue-like elasticity, which may offer an additional advantage for myogenesis.

### 2.7 Implant operation and muscle regeneration *in vivo*

To demonstrate the potential for grafting within a living organism, muscle scaffolds were implanted as volumetric constructs to regenerate previously injured tibialis anterior (TA) muscle in mice. Figure 11 shows the surgical procedure that was used to implant the 10-day matured scaffolds as volumetric TA muscle models. A volumetric muscle deficit was induced in the right leg of the mouse by surgically removing approximately 30% of the mass of the TA muscle. The developed muscle scaffold, measuring approximately ∼2.5 mm in diameter and ∼6 mm in length, was positioned within the defect area. It was then secured to the host muscle using 6-0 black silk sutures. Subsequently, the fascia and skin were sutured using 4-0 black silk stitches to complete the closure. The muscle-implanted mice were conscious and could access food and water 2 hours after surgery. They exhibited healthy activities, no fever, and no infection for 4 weeks after implantation (Videos S7-10). The defect (injured, no scaffold), Mo-M, and Mu-C mice had a limp 1 day after implantation compared to the control mouse (Video S7). Only the defect mouse still had limp 1 week after implantation (Video S8). The Mu-C mouse walked normally 2 weeks after implantation and moved sprightly after 4 weeks compared with the control mouse (Videos S9 and 10). These results demonstrated that the Mu-C mouse recovered quickly from the TA muscle defect.

**Figure 11:**
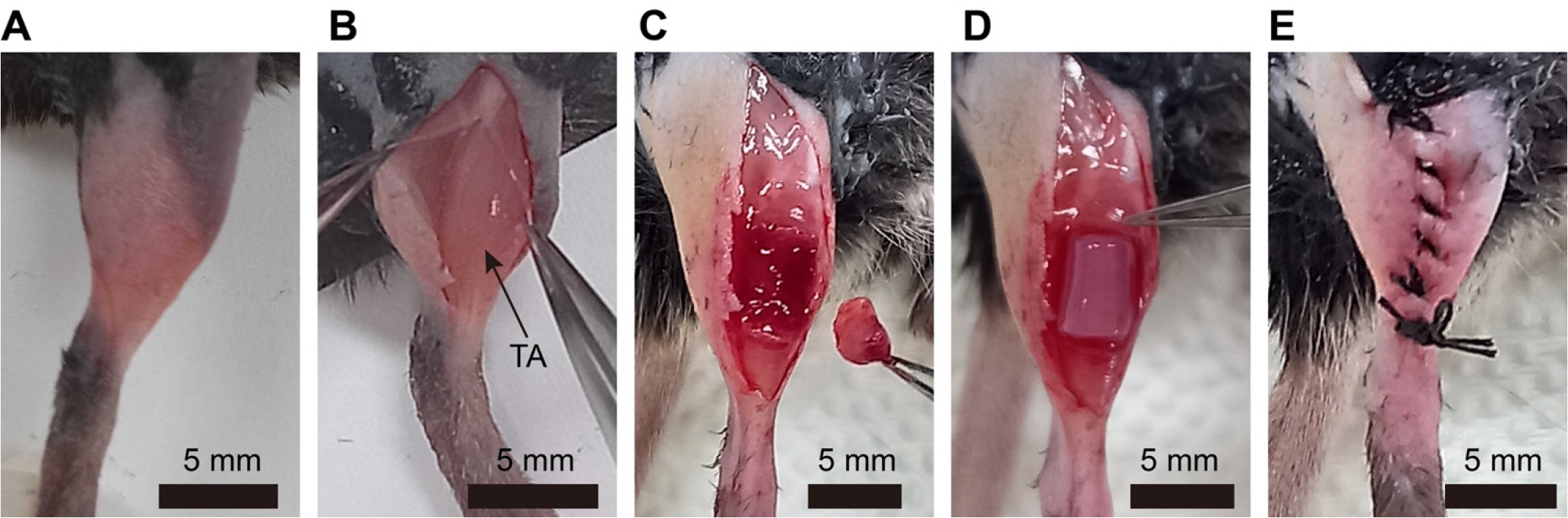
The process of implantation surgery in a mouse. (a) After anaesthesia, the hair on the thigh skin of the mouse was removed. Povidone-iodine (10%) was applied to the area for disinfection before implantation surgery. (b) An incision in the thigh skin was made using sterilized scissors and fine forceps. The size of the opened area was approximately 20 mm long and 15 mm wide, and the TA muscle was visible. (c) Approximately 30% of the TA muscle was excised to create a volumetric defect. (d) The fabricated muscle bundle (approximately 2.5 mm in diameter, 6 mm long) was inserted into the defective region and stitched to the host muscle using black silk 6-0 sutures. (e) The fascia and skin were closed using black silk 4-0 sutures.

Four distinct in vivo conditions were established to compare the effect of volumetric muscle implantation in TA muscle regeneration, including a defect (injured, no scaffold), a control group with no implantation, serving as a baseline for comparison with mice implanted with Mo-M and Mu-C scaffolds (Figure 12a). The gross morphology of the harvested TA muscles from the defect mouse showed permanent muscle loss (Figure 12b). Moreover, the TA muscle volume of the scaffold-implanted groups increased 4 weeks after implantation (Figure 12b). Figures 12c and d show histological images stained with haematoxylin–eosin (HE) and Masson’s trichrome 4 weeks after implantation. The diameter and shape of the individual muscle fibres of the Mu-C scaffold-implanted mouse were visibly similar to those of the control mouse (Figure 12, Figure 13a). There were many more wavy muscle fibres in the defect mouse and the Mo-M scaffold-implanted mouse than in the Mu-C scaffold-implanted mouse (Figure 12, Figure 13b). Furthermore, increased collagen deposition was detected in the defect mouse and the Mo-M scaffold-implanted mouse compared to the Mu-C scaffold-implanted mouse (Figure 12d, Figure 13c). These results indicated that there was a lower tendency of scar tissue formation in the Mu-C scaffold-implanted mouse than in the defect and Mo-M scaffold-implanted mice [41, 42]. Video S11 shows the contraction of the muscle bundle dissected from the Mu-C scaffold-implanted mouse 4 weeks after implantation.

**Figure 12:**
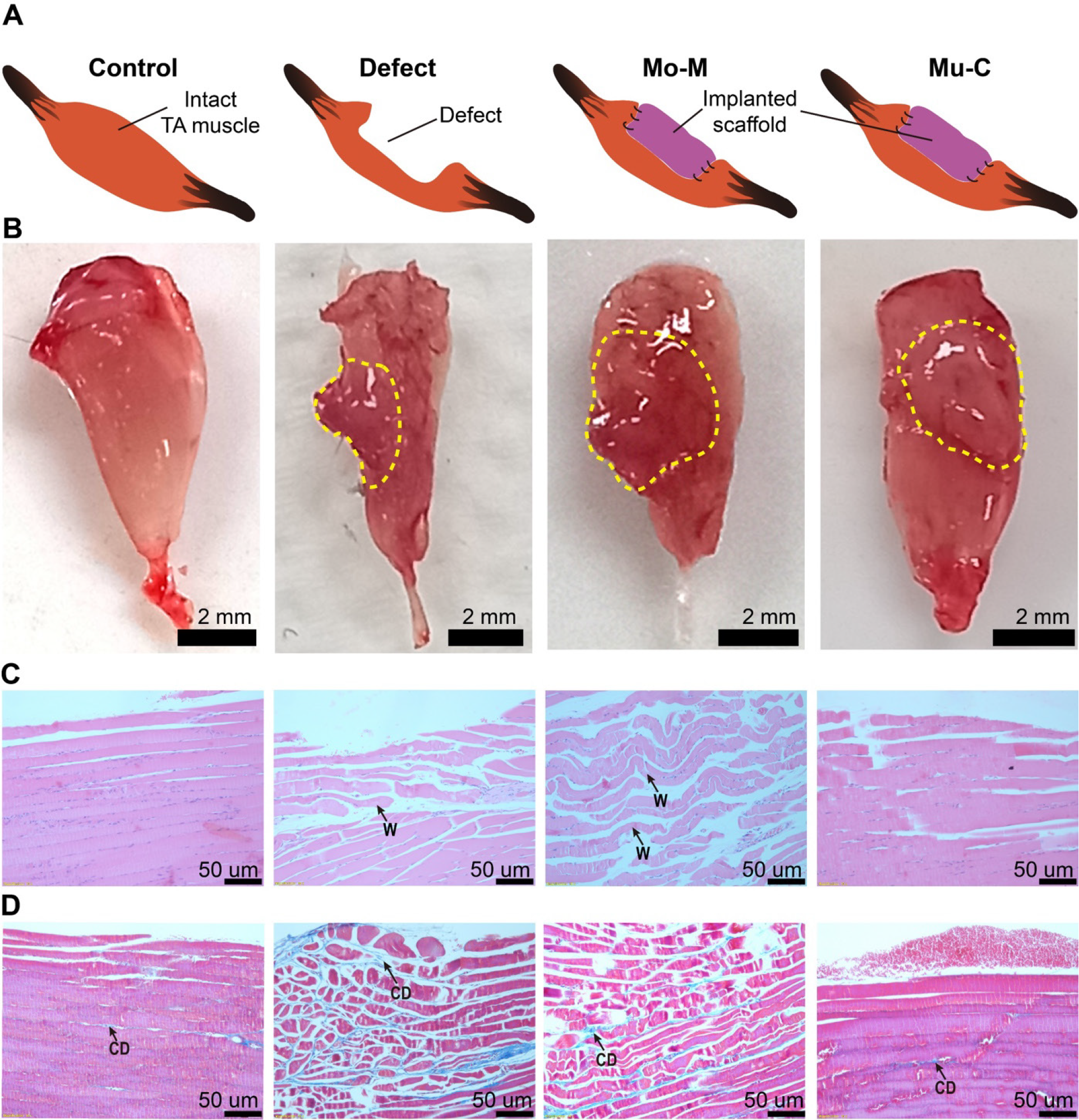
Extracted muscles 28 days after implantation. (a) Conceptual schematics of the control, defect, Mo-M implantation, and Mu-C implantation conditions. (b) Extracted TA muscle samples of the respective groups 28 days after implantation. (c) Histological images of representative extracted muscles from the four groups stained with HE 28 days after implantation. The wavy muscle fibres are marked as “W”. (d) Histological images of the extracted muscles stained with Masson’s trichrome (MTS) 28 days after implantation. Collagen deposition is marked as “CD” (blue spots).

**Figure 13:**
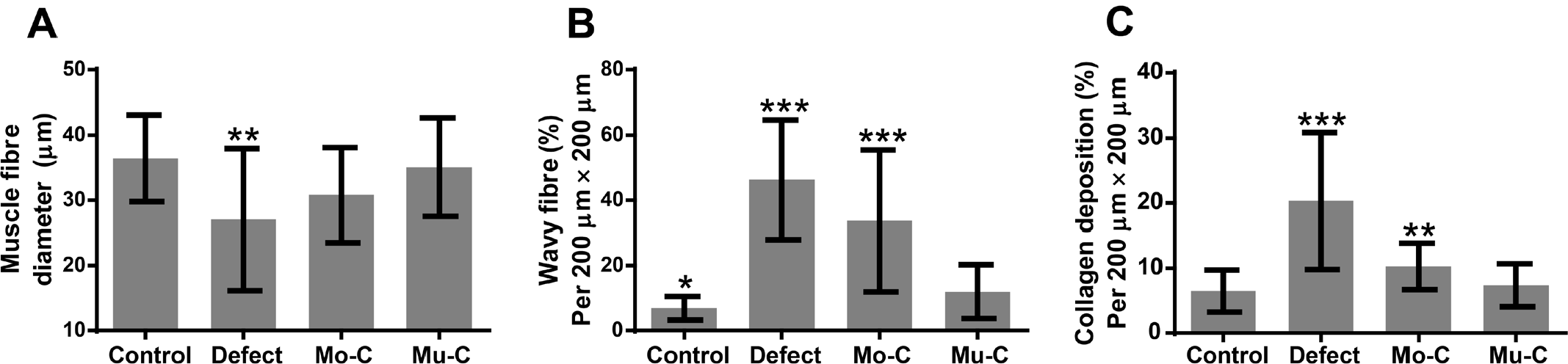
Quantitative analysis of the histological images from the control, defect, Mo-M implantation, and Mu-C groups. (a) Muscle fibre diameter (** 0.001 < p < 0.01 comparing the Mu-C group to the other groups; n ≥ 20). (b) Number of wavy muscle fibres in 200 µm × 200 µm cross-sections (* 0.01 < p < 0.05, ***p < 0.001 comparing the Mu-C group to the other groups; n ≥ 20). (c) Collagen deposition in 200 µm × 200 µm cross-sections (* 0.01 < p < 0.05, ***p < 0.001 comparing the Mu-C group to the other groups; n ≥ 20).

## 3. CONCLUSION

Here, we developed a method to fabricate multifascicle human skeletal muscle scaffolds that mimic the natural structure of human skeletal muscle bundles using a seven-barrel nozzle combining with co-culturing of HUVECs, HFs, and hSkMDCs. The muscle scaffold exhibited a favourable microenvironment to promote cell proliferation, differentiation, vascularization, mechanical properties, and functionality.

Furthermore, in an in vivo mouse model of volumetric muscle loss, the muscle scaffold effectively regenerated the tibialis anterior muscle defect, demonstrating its potential for volumetric muscle transplantation. The multi-barrel nozzle apparatus was employed to generate functional Mu-C muscle scaffolds that closely replicated the structural characteristics of human skeletal muscle. Our nozzle would also be further used to produce other volumetric functional tissues, such as tendons, peripheral nerves, and liver tissue.

